# Structure-based inference of eukaryotic complexity in Asgard archaea

**DOI:** 10.1101/2024.07.03.601958

**Authors:** Stephan Köstlbacher, Jolien J. E. van Hooff, Kassiani Panagiotou, Daniel Tamarit, Valerie De Anda, Kathryn E. Appler, Brett J. Baker, Thijs J. G. Ettema

## Abstract

Asgard archaea played a key role in the origin of the eukaryotic cell. While previous studies found that Asgard genomes encode diverse eukaryotic signature proteins (ESPs), representing homologs of proteins that play important roles in the complex organization of eukaryotic cells, the cellular characteristics and complexity of the Asgard archaeal ancestor of eukaryotes remain unclear. Here, we used *de novo* protein structure modeling and sensitive sequence similarity detection algorithms within an expanded Asgard archaeal genomic dataset to build a structural catalogue of the Asgard archaeal pangenome and identify 908 new ‘isomorphic’ ESPs (iESPs), representing clusters of protein structures most similar to eukaryotic proteins and that likely underwent extensive sequence divergence. While most previously identified ESPs were involved in cellular processes and signaling, iESPs are enriched in information storage and processing functions, with several being potentially implicated in facilitating cellular complexity. By expanding the complement of eukaryotic proteins in Asgard archaea, this study indicates that the archaeal ancestor of eukaryotes was more complex than previously assumed.

## Introduction

The origin of the eukaryotic cell, with its complex and compartmentalized features, is regarded as the biggest evolutionary discontinuity since the advent of cellular life on Earth (*1*). Yet, many key details regarding eukaryogenesis (the series of evolutionary events that lead to the emergence of the eukaryotic cell from prokaryotic ancestors some 2 billion years ago (*2, 3*), remain elusive. The eukaryotic cell is the result of a symbiosis comprising an archaea-related host cell (*4, 5*) and a bacterial endosymbiont, the mitochondrial progenitor (*6, 7*). While the identity of the endosymbiont was traced back to the Alphaproteobacteria several centuries ago (*8, 9*), the archaeal host remained obscure until recently. This changed with the discovery of Asgard archaea, which were shown to represent the closest prokaryotic relatives of the archaeal host cell from which eukaryotes evolved (*10*–*13*). Analysis of Asgard archaeal genomes revealed the presence of numerous homologs of proteins previously deemed eukaryote-specific – so-called Eukaryotic Signature Proteins (ESPs) (*14*). Intriguingly, many of these ESPs represent building blocks fundamental for eukaryotic cellular complexity, including proteins essential to vesicular biogenesis and trafficking, and to the dynamic eukaryotic cytoskeleton. Recent work has indicated that several Asgard ESPs indeed represent functionally equivalent homologs of eukaryotic proteins (*15*–*18*), suggesting that Asgard archaea might display eukaryote-like cellular features beyond the dynamic actin cytoskeleton observed in the first enrichment cultures (*19, 20*). However, the detailed cellular characteristics and level of complexity of present-day Asgard archaea and of the Asgard archaeal ancestor of eukaryotes remain unclear.

In addition to enabling making inferences about cell-biological properties and the lifestyle of present-day Asgard archaeal lineages, identifying and characterizing ESPs aids in reconstructing the ancestral Asgard lineage from which eukaryotes evolved. Yet, the identification process is currently limited by several factors. First, the definition of ESPs has proven challenging as increasingly sensitive homology search algorithms and improved sampling of genomic diversity across the tree of life have facilitated the discovery of ESP homologs in diverse prokaryotes (*11, 13, 21*), including Asgard archaea (*10, 11, 13, 21*). Although this has increased the fraction of proteins with a prokaryotic provenance in the last eukaryotic common ancestor (LECA), it has also been steadily decreasing the number of *sensu stricto* ESPs (proteins unique to eukaryotes). Therefore, a more relaxed definition of ESPs has been adopted, referring to proteins associated with conserved key eukaryotic processes (*5*), or more specifically related to cellular complexity (*20*). However, such a function-centered definition is problematic since many eukaryotic proteins remain poorly characterized, in particular if they are absent in model organisms such as yeast and human, yet could potentially play key roles in fundamental eukaryotic processes. Another confounding factor in identifying ESPs of prokaryotic origin involves the limits of reliable sequence homology detection. As sequence similarity decreases, it becomes increasingly challenging to infer homology between two proteins (*22*). The stem separating eukaryotes from their archaeal relatives represents one of the longest branches in the tree of life (*12, 13*). Hence, sequences from present-day Asgard archaea and eukaryotes have diverged extensively, and homology might not even be reliably detected, even when using sensitive methods (*22*). However, protein structure is several times more conserved than protein sequence (*23*), and structural information has been shown to increase sensitivity of sequence homology inference (*24*). Recent advances in *de novo* protein structure prediction using AlphaFold (*25*) and related tools enable the large-scale generation of high-quality protein structure models. Combined with new methods to efficiently search large databases for similar structures (*26*), it has become feasible to identify highly diverhent homologs by using structural information (*27, 28*).

Here, we explore these recent advances in protein structure prediction and comparison tools to expand the identification and characterization of ESPs in Asgard archaea beyond sequence similarity. By analyzing an extended Asgard archaeal pangenome, we identified 908 new structure-based ‘isomorphic’ ESPs (iESPs), more than tripling the overall number of reported Asgard ESPs. Our structural catalogue of the Asgard archaeal pangenome reveals a marked increase of Asgard ESPs involved in information storage and processing, and in cellular processes and signaling, suggesting that the archaeal ancestor of eukaryotes was more eukaryote-like than was previously assumed.

### Structural modeling of the Asgard archaeal pangenome

To generate structural models of representative proteins encoded by the Asgard archaeal pangenome, we analyzed a diverse set comprising 936 Asgard archaeal draft genomes (Fig. 1A, Data S1), including 404 metagenome-assembled genomes (MAGs) that were obtained in a recent study (*29*). In addition to the previously sampled Asgard archaeal diversity (*13, 30*), this expanded dataset encompasses MAGs from Atabeyarchaeia (*31*) and Ranarchaeia (*29*), two additional deep-branching clades (Fig. S1A). We grouped protein sequences encoded by these Asgard genomes by combining reference-based clustering into previously established Asgard clusters of orthologous genes (AsCOGs) (*21*) with *de novo* gene clustering (Fig. 1B). This resulted in 96% of Asgard archaeal proteins grouped in 37,313 clusters of at least five proteins, including 22,609 *de novo* clusters (Fig. 1B). For computational feasibility, we selected one evolutionary representative protein sequence per cluster (see *Methods*) to generate a high-quality structural model (Fig. 1C).

**Fig. 1.**
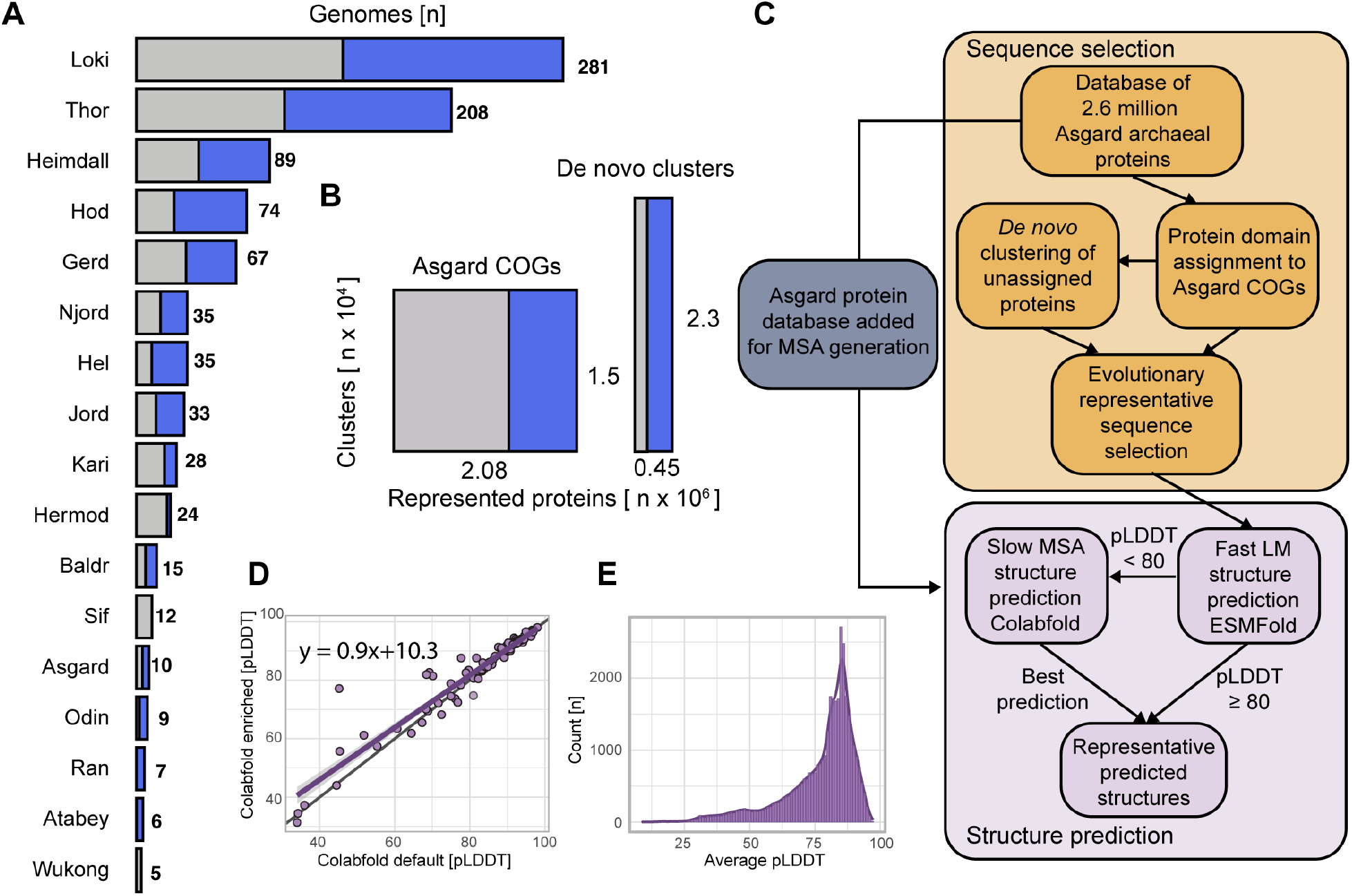
Modeling the Asgard archaeal structural pangenome. (**A**) Number of Asgard archaeal draft genomes per group in the database used for pangenome-wide structural analyses. Fill color indicates publicly available genomes (grey) and newly added Asgard archaeal draft genomes (blue), respectively. (**B**) Protein sequence clustering into existing Asgard COGs and de novo clustering with unassigned proteins. X-axis indicates the number of proteins and y-axis the number of respective clusters. Fill indicates protein sequences from publicly available genomes (grey) and added Asgard archaeal draft genomes (blue), respectively. (**C**) Workflow for the pangenome-wide prediction of Asgard archaeal protein structures. (**D**) Scatter plot depicting pLDDT scores of structure predictions of 100 randomly selected ‘*Candidatus* Prometheoarchaeum syntrophicum’ proteins computed with the default (x-axis) and the Asgard-enriched (y-axis) ColabFold database, respectively. The diagonal black line indicates x = y, purple line indicates linear correlation fitted to the data. (**E**) Distribution of average pLDDT scores of 37,223 predicted Asgard archaeal protein structures. MSA, multiple sequence alignment.

To determine an efficient and effective approach for *de novo* structure prediction, we modelled structures for 100 randomly selected proteins of the Asgard archaeon ‘*Candidatus* Prometheoarchaeum syntrophicum’ (Data S2). As AlphaFold relies on homology information to predict protein structure, it tends to perform poorly if few homologs are found within its reference sequence database (*25*). We therefore used ColabFold (*32*), an accelerated AlphaFold workflow, and expanded the database with all available Asgard protein sequences. In addition, we used ESMfold (*33*), a prediction tool based on a protein language model (pLM) that circumvents the time-consuming sequence homology search. We classified predictions as high-quality if they had an average predicted local distance difference test (pLDDT) score of at least 80. We found that incorporating the Asgard proteins to the ColabFold homology search database led to better models for some proteins (Fig. 1D, Fig. S1B). Overall, we obtained the most high-quality structure predictions when combining pLM and sequence alignment-based techniques (Fig. S1C). We decided to predict structures for each representative protein sequence using the fast ESMfold algorithm, and only if its average pLDDT score was below 80, we used the more time-consuming ColabFold method (Fig. 1C and S1C-D). This approach resulted in 37,223 predicted structures with a median pLDDT of 82 (interquartile range 71-86), covering 99.8% of all clusters (Fig. 1E).

### Structures facilitate sequence annotation beyond the twilight zone of sequence similarity

Next, we aimed to annotate the protein clusters by identifying homologs, using sensitive multiple sequence alignment (MSA) methods and their representatives’ predicted Asgard protein structures (Fig. 2A). Using traditional MSA-based searches, we obtained high-confidence hits (HHsearch P≥95) to the COG/KOG database for 29% (n=10,681) of the protein clusters. With structure-based similarity searches, we retrieved significant hits in the SwissProt database for 47% (n=17,309) of representative proteins (Fig. 2B). We could annotate 8,891 proteins with both sequence and structure, finding agreement of the COG assignments for 96% of representative proteins and their respective best structural hits in SwissProt. Of note, almost half of the protein representatives with both a highly confident (sequence-based) COG and structural hit displayed less than 20% sequence identity to their best structure hit (n=4,263; median of 18.6%; interquartile range (IQR) =14.2-28.0%), falling below the ‘twilight zone’ of sequence identity (the zone between 20–35% sequence identity where homology becomes challenging to predict with regular algorithms) (*22*). This demonstrates the high sensitivity of MSA-based searches. We found that all protein representatives that could only be annotated with MSA-based searches did have a structure hit in UniProt50 but belonged to protein families not annotated in SwissProt. To illustrate the ability of our approach to annotate protein clusters even in cases of low sequence identity, we recovered the recently discovered distant Asgard archaeal homolog of Vps29 (*13*), a component of the eukaryotic retromer and retriever complexes, with sequence similarity searches (best structure hit amino-acid identity=27.5%; HHsearch P=99.8), as well as with local and global structural alignment (Foldseek E-value=1.9·E^-20^, DaliLite Z-score= 30, Fig. 2C).

**Fig. 2.**
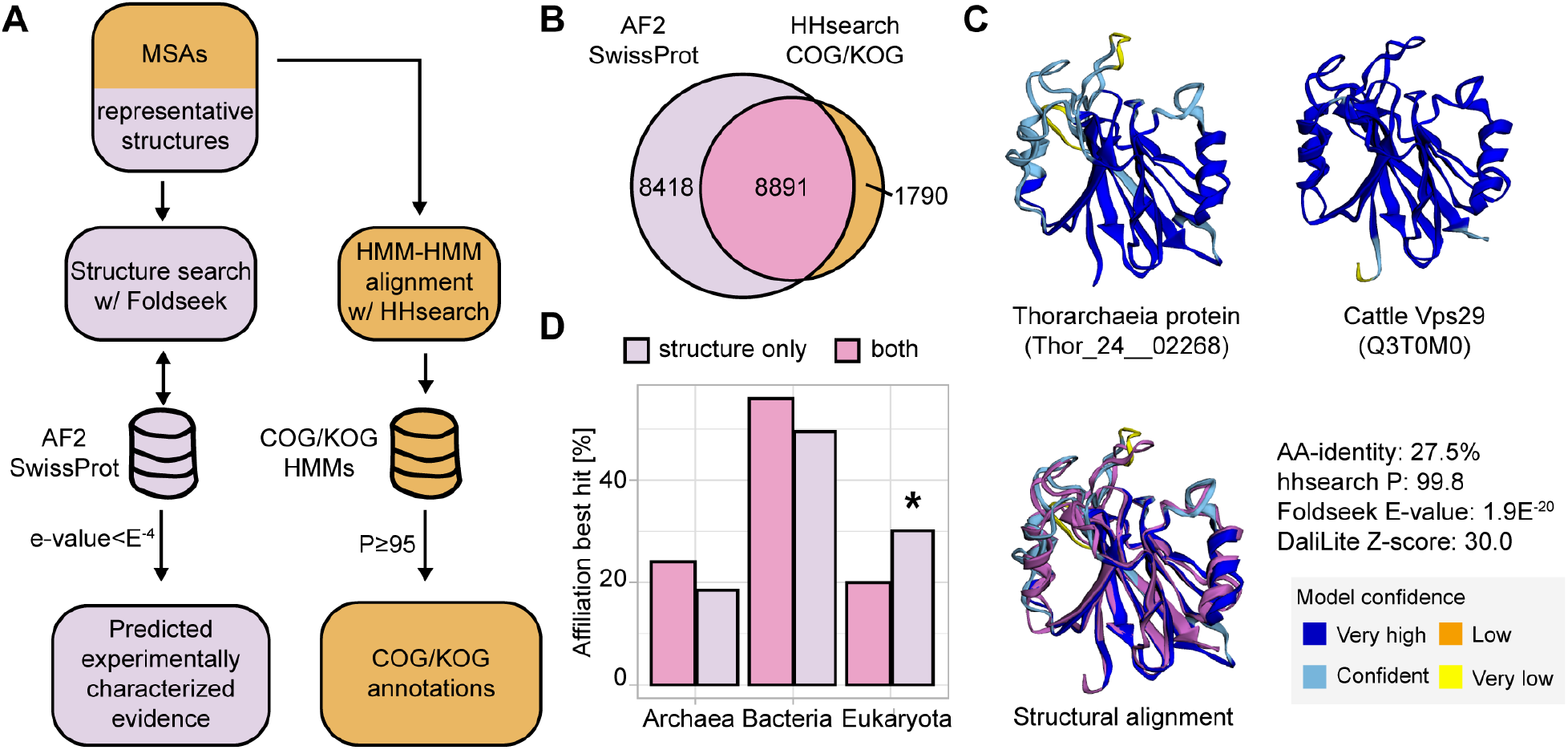
Structural information recovers significantly more eukaryotic best hits. (**A**) Workflow to annotate Asgard archaeal proteins based on homology using sequence and structural similarity. (**B**) Venn diagram depicting the number of clusters or cluster representing protein structures annotated using HHsearch against the COG/KOG database (orange) and structural searches against AF2 SwissProt (violet), respectively. The intersection of both techniques is marked in pink. (**C**) Structure prediction of Vps29 Asgard archaeal representative (left), its most similar SwissProt prediction (right; Cattle Vps29, Q3T0M0), and their overlay with the eukaryotic protein in violet (bottom). HMM, hidden Markov model; AA-identity, amino acid identity to best structure hit; P, hhsearch probability. (**D**) Bar plot depicting the proportion of the domain-rank affiliation of best SwissProt structure hit based on annotation with both structural and sequence assignment (pink), or just structure (violet). The structure-only annotation is significantly enriched in eukaryotic best hits as indicated by the asterisk (one-tailed Fisher’s exact test p-value: 1.9·E^-63^).

Subsequently, we tested whether differences in taxonomic assignment existed for proteins that could only be annotated based on structure, as these likely exhibit stronger sequence divergence. The 8,418 proteins exclusively annotated using structures showed significantly lower sequence identities to their best structure hits (medians=15.1% vs 18.6%; IQR=12.0-20.2% vs 14.2-28.0%, Wilcoxon signed-rank test p-value: 5·E^-16^; Fig. S2) and were, interestingly, enriched in best hits against eukaryotic protein structures (Fig. 2D). This could indicate that, for Asgard archaeal proteins, their eukaryotic homologs diverged more extensively compared to their prokaryotic homologs. We further observed that these proteins are enriched in functional categories related to cellular processes and signaling (Bonferroni-corrected one-tailed Fisher’s exact test p-value: 9.3·E^-77^), and more specifically in “intracellular trafficking, secretion, vesicular transport”, “signal transduction”, and “extracellular structures” (Bonferroni-corrected one-tailed Fisher’s exact test p-values: 5·E^-6^, 3·E^-4^, 6.3·E^-4^, respectively) (Fig. S2).

### Asgard archaeal protein structures isomorphic to eukaryotic proteins

Next, we used structure-based similarity searches to identify novel ‘isomorphic’ ESPs in Asgard archaea (Fig. 3A), hereafter referred to as iESPs. We define an iESP as an Asgard archaeal protein structure that exhibits a statistically significant overrepresentation of eukaryotic protein structures in (*i*) all hits or (*ii*) the top 95% bit-score quantile of hits (Fig. 3B; *Methods*). We identified 1,319 iESPs that have thus far not been identified as Asgard archaeal ESPs (Fig. 3B). Of note, we only captured 46% (611 proteins) of the 1,323 previously established Asgard archaeal ESPs, indicating that previous definitions for ESPs have been rather permissive (also see above; Fig. 3B; Data S3). For example, 40 AsCOGs containing roadblock domains are considered ESPs and Asgard archaeal proteins have been shown to form similar structures to their eukaryotic relatives (*34*). However, only four (cog.000673, cog.000921, cog.006948, cog.008459) are enriched in eukaryotes according to our representative structures. Indeed, roadblock/LC7 domain (PF03259) containing proteins are common in prokaryotes with 24,892 and 2,494 such proteins encoded by bacterial and archaeal genomes, respectively, compared to 5,724 proteins in eukaryotes (Pfam database accessed 12^th^ June 2024).

**Fig. 3.**
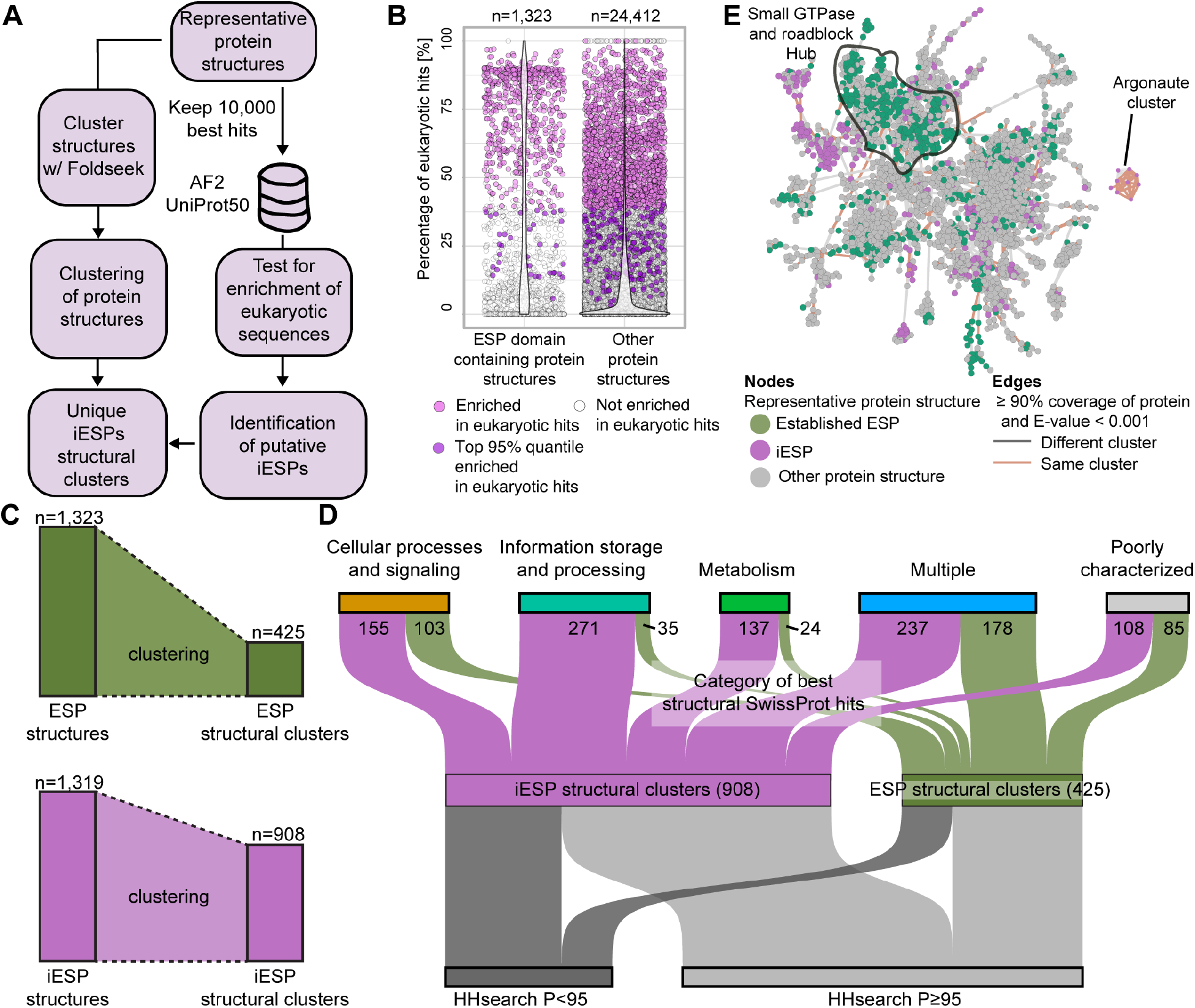
Structure-guided identification of functionally diverse iESP structural clusters. (**A**) Workflow to cluster protein structures and identify iESPs. (**B**) Identification of Asgard archaeal iESPs based on structural similarity. (**C**) Bar chart summarizing the clustering of previously described ESP and iESP protein structures into structural clusters, respectively. (**D**) Sankey diagram displaying functional categories of newly identified iESPs clusters and clusters containing previously established ESPs. Categories are inferred from the best SwissProt hits EggNOG annotation. ‘Multiple’ indicates an association of a structural cluster with multiple functional categories. (**E**) Subgraph of protein structure similarity network, highlighting small GTPase (black outline) and Argonaute proteins. P, probability.

To reduce redundancy, and to obtain an overview of the structural connectivity within the (i)ESP landscape, we clustered the 37,223 predicted Asgard archaeal protein structures based on their similarity, which we delineated into 19,775 structural clusters (see *Methods* and Fig. 3A and S3A). In total, the 1,319 newly identified iESPs and all 1,323 previously identified ESP protein structures are contained in 908 and 425 clusters (Fig. 3C), respectively, indicating that our structure-based approach more than triples the potential number of Asgard archaeal proteins that entered the eukaryotic stem lineage. A high-level functional assessment revealed remarkable differences between iESP and ESP structural clusters (Fig. 3D Data S3). For example, 64% of previously identified ESP clusters (336 of 425) have functions in cellular processing and signaling, including a hub of 59 clusters collectively encompassing 932 Asgard archaeal small GTPase protein representative structures (Fig. 3E), which are known to have undergone extensive duplication in both eukaryotes and Asgard archaea (*10, 11, 21, 35, 36*). In contrast, only 28% of iESP clusters (258 of 908) are involved in cellular processing and signaling functional (when including clusters containing multiple functional categories). Among these, we identified a single cluster containing eight Argonaute-related Asgard archaeal iESPs (Fig. S3). Argonautes are involved in DNA and RNA interference in prokaryotes and eukaryotes, respectively (*37*). A recent study indicated that some Asgard archaeal Argonautes appear to be functionally related to their eukaryotic counterparts (*38, 39*). We obtained best structural hits to eukaryotic AGO and PIWI proteins (Fig. 3E and S3), illustrating their stringent structural conservation despite their high level of sequence divergence (*37*).

We also retrieved many iESP clusters specific to metabolism (Fig. 3D n=137), which was thus far poorly represented among previously found ESPs in Asgard archaea (n=24). For example, we identified diverse iESPs, including best hits to proteins of the eukaryote-type mevalonate pathway (phosphomevalonate kinase, Swissprot accession: Q2KIU2), the oxygen-dependent degradation of prenylated proteins (PCYOX1, Q5R748), and reactive oxygen species defense (SOD1, P80566). As an outstanding feature, we identified many iESP clusters involved in information storage and processing functions (n=271), of which 169 are related to translation, ribosomal structure and biogenesis, a function in eukaryotes that is known to have an archaeal provenance (*40*). iESPs identified within the latter functional category included best structural hits to eukaryotic elongation factor 1A lysine methyltransferase 1 (EEF1AKMT1, Q17QF2) and the malignant T-cell-amplified sequence 1 that is involved in translation re-initiation (MCT-1, Q2KIE4) (Data S3). Altogether, our structure-based and functionally unbiased approach identified hundreds of new ESPs, bearing relevance for efforts to reconstruct the physiology and cell biological features of both extant Asgard archaea as well as the archaeal ancestor or eukaryotes.

### Asgard archaeal iESPs potentially implicated in cellular complexity

The emergence of intricate cellular compartments has been a hallmark process of eukaryogenesis, yet the origins of many genes responsible for the formation of these compartments remain elusive (*41*). To identify Asgard archaeal proteins potentially involved in cellular compartmentation, we investigated iESPs with robust structural assignment but limited, ‘twilight zone’ sequence similarity (Fig. 3D) and examined their relationship to their evolutionary eukaryotic counterparts. By employing targeted sequence-based searches with iterative refinement guided by structural similarity, we managed to link several iESPs at the sequence level, after which we constructed multiple sequence alignments and performed phylogenetic analyses (see *Methods*).

One of the eukaryotic complexes with a role in cell compartment biology and lacking a clear prokaryotic ancestry is the vault, the largest reported ribonucleoprotein complex conserved in diverse eukaryotes and suggested to be involved in transport between cellular compartments, signal transmission, cellular stress protection, and immune response (*42*). Vaults are primarily composed of two symmetric cups, each consisting of 39 molecules of the major vault protein (MVP) (*43*). While prokaryotic homologs of MVP have so far only been described in a few Bacteria (*44*), we identified an Asgard archaeal protein structure with a reciprocal best hit to *Xenopus laevis* MVP (Q6PF69, Fig. S4). In total, we found ten Asgard archaeal MVP homologs, half of which in our phylogenetic analysis affiliate with a clade including eukaryotic MVPs (Fig. 4A and S4A). The representative Asgard archaeal MVP displays a structure similar to the resolved rat MVP, including the cap helix, shoulder, and repeat domains, even though the Asgard archaeal homolog only contains five instead of nine repeat domains present in the rat protein (*45*) (Fig. 4B). Multimer structure modeling suggests a closed cup with 10 Asgard archaeal MVP molecules (interface predicted template modelling score, ipTM=0.525, average pLDDT=71.4, Fig, S4B-C) that is markedly smaller than the eukaryotic representative, which displays 39 MVP molecules (Fig. 4C) (*45*). While the role of MVP homologs in Asgard archaea remains unknown, our findings support a prokaryotic, and possibly Asgard archaeal origin of the eukaryotic MVP.

**Fig. 4:**
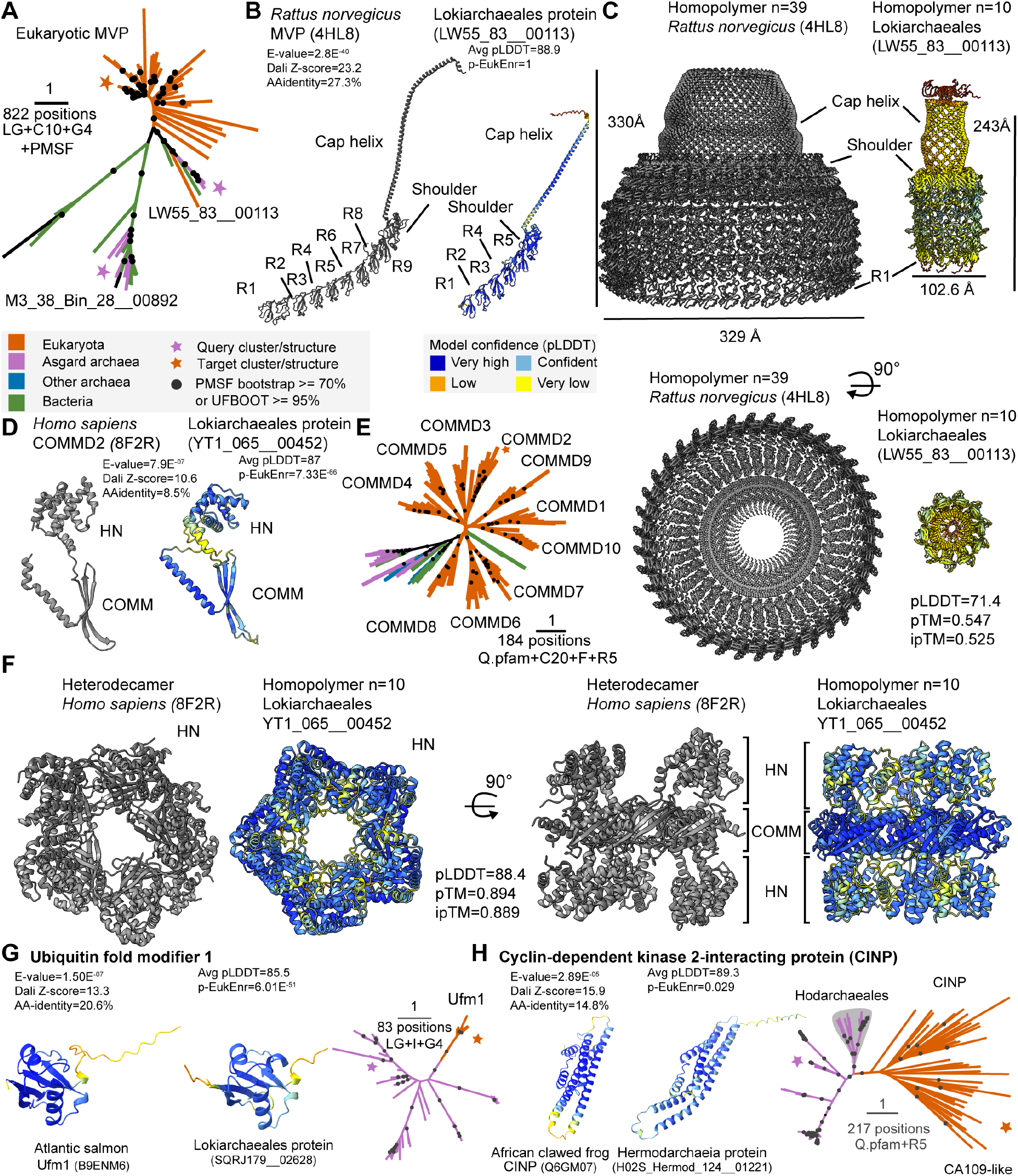
Asgard archaeal protein complexes implicating cellular compartmentalization. Asgard archaeal proteins related to eukaryotic (**A-C**) MVPs and (**D-F**) COMMD-containing proteins. (**A**) Phylogeny of prokaryotic and eukaryotic full-length MVPs. See Fig. S4A for tree based only on the shoulder domain. (**B**) Rat MVP complex (*45*) next to Lokiarchaeial MVP (predicted structure) indicating the cap helix, shoulder, and repeat domains (R). (**C**) Biological assembly of the rat MVP cap (left) next to a multimer model of the Asgard archaeal homodecamer (right). (**D**) Human COMMD2 next to Lokiarchaeial homolog indicating the HN and COMM domains. (**E**) Phylogeny of prokaryotic and eukaryotic COMMD-containing proteins. (**F**) Resolved human COMMD heterodecamer (*46*) next to a multimer model of the Asgard archaeal homodecamer. (**G, H**) Identification of Asgard archaeal iESPs of eukaryotic ubiquitin fold modifier 1 (**G**) and cyclin-dependent kinase 2-interacting protein (Hodarchaeales clade indicated with grey background) (**H**). Asgard archaeal query protein structure, best-scoring SwissProt target structural model and phylogenetic analysis of related protein sequences are indicated in the left, middle and right panel, respectively. Structural models exclude long terminal disordered regions. Additional data include Foldseek E-value, Dali Z-score, enrichment of eukaryotic structures (Fisher’s exact test, Bonferroni-corrected p-value, ‘p-EukEnr’), and amino-acid identity to best structure hit (‘AA-identity’). Phylogenetic analyses highlight sequences for query and target structures, input MSA positions, and substitution model. Scale bar: 1 amino acid substitution per position. Multimer model confidence measures (pLDDT, pTM, ipTM) are indicated.

Another eukaryotic complex with an elusive origin is Commander, which is required for endosomal recycling of diverse transmembrane cargos and is composed of sixteen subunits that are arranged into the CCC and retriever subcomplexes. While some retriever components have been reported in Asgard archaea before (Vps29, Fig. 2C; Vps35) (*46*), the CCC (named after its components CCDC22, CCDC93 and COMMD) subunits, including the heterodecamer-forming COMMD proteins, thus far lacked prokaryotic homologs (*46*). Our structure-based searches retrieved an Asgard archaeal iESP that displayed the characteristic COMMD protein structure, i.e., an α-helical N-terminal (HN) and a C-terminal COMMD domain (*47*), while displaying extremely low sequence identity (8.5%) (Fig. 4D). Subsequent sensitive HMM-based searches yielded homologs in diverse Asgard archaea (Lokiarchaeales, Helarchaeales, and Heimdallarchaeia), and some other prokaryotes. In our phylogenetic analysis, eukaryotic COMMD proteins (COMMD1-10) form a near-monophyletic group (Fig. 4E), confirming that eukaryote-specific gene duplications gave rise to the COMMD heterodecamer (*46, 48*). While our phylogenetic analyses failed to resolve the origin of eukaryotic COMMD, multimer modeling of an Asgard archaeal homolog suggests that eight, ten, or 12 molecules may form a homomultimeric complex with high confidence (n=10; ipTM=0.889, pLDDT=88.4; Fig. 4F, Fig. S4D-E).

Among the identified iESPs, Ubiquitin fold modifier 1 (Ufm1) has previously not been reported outside of eukaryotes. Despite limited sequence similarity, Ufm1 exhibits structural similarities to ubiquitin (*49*) and is implicated in DNA damage and ER stress responses, although it has not been characterized extensively (*50*). We identified Ufm1 homologs in nine of the major Asgard archaeal clades, but not in any other prokaryote (Fig. 4G), indicating an Asgard archaeal provenance of Ufm1 in eukaryotes. Similarly, no prokaryotic homologs have yet been reported for the cyclin-dependent kinase 2-interacting protein (CINP), a protein involved in DNA replication complex and DNA damage control (*51, 52*) that was recently also implicated in eukaryotic ribosome biogenesis (*53*). Our sequence similarity searches revealed it is present in five major Asgard clades, but not in other prokaryotes. Phylogenetic analyses revealed that eukaryotic sequences are monophyletic and cluster with Hodarchaeal sequences with good support (Fig. 4H, UFBOOT: 99%), suggesting that eukaryotes inherited this protein from their Hodarchaeal ancestor (*13*).

## Discussion

Large scale analyses of the protein structure universe are becoming powerful approaches to identify origins and functions of proteins beyond the capabilities of standard sequence-based homology searches (*54, 55*). Here, we explored the development of these tools to gain insight into the archaeal provenance of the eukaryotic cell. By building and analyzing a structural catalogue of the Asgard archaeal pangenome, we improved the annotation of Asgard archaeal proteins lacking significant sequence similarity. Our approach revealed many Asgard archaeal protein families, iESPs, that are structurally most similar to those of eukaryotes. As in previous studies that relied on sequence similarity searches to identify ESPs (*10, 11, 13, 21*), we also identified iESPs involved in cellular processes and signaling, including many that participate in intracellular trafficking, secretion and vesicular transport. However, our extended analyses retrieved many iESPs involved in additional processes, such as information storage and processing. This observation is in line with the general conception that many eukaryotic proteins involved in translation, transcription, replication and DNA repair have an archaeal provenance (*56*). Furthermore, we found that iESPs are also relatively enriched in metabolic functions, which contrasts with previous work indicating that metabolic functions in eukaryotes predominantly are of bacterial origin (*57, 58*). The underlying reason for this observation is unclear. Yet, more likely, and in congruence with recent work showing that eukaryotic central carbon metabolic pathways are in part of Asgard archaeal origin (*59*), these metabolic iESPs represent ancient homologs of eukaryotic proteins that have evolved beyond the limit of reliable sequence similarity detection. Altogether, our analyses suggest that a thus far underappreciated fraction of the eukaryotic metabolic repertoire is of Asgard archaeal provenance.

While several studies have revealed that some ESPs, such as small GTPases, actin homologs and several subunits of the ESCRT complex, are nearly universally distributed across Asgard archaeal genomes, many ESPs display a rather patchy distribution (*11, 13, 21*). This patchiness is evident, for example, for Asgard archaeal homologs of adaptor proteins, Golgi-associated retrograde protein (GARP), homotypic fusion and protein sorting (HOPS) and class C core vacuole/endosome tethering (CORVET) complexes (*13*). A similar observation can be made for iESPs, which predominantly display patchy distribution patterns across Asgard archaeal taxa. These patchily distributed ESPs and iESPs likely represent ancient protein families that were already present in the Asgard archaeal lineage from which eukaryotes emerged, and have bene subjected to multiple loss events or horizontal gene transfers among Asgard archaeal lineages. Overall, given their patchy distribution, combined with the evolutionary distance between present-day Asgard archaeal and eukaryotic proteins, it remains unclear to what extent Asgard archaeal iESPs are functionally equivalent to their eukaryotic counterparts. While structural conservation has been shown to be tightly linked to protein function, even at high levels of sequence divergence (*60*), future studies are needed to corroborate the functions of Asgard archaeal iESPs and ESPs. Such studies, complemented with cultivation efforts, are ultimately needed to elucidate the biology of Asgard archaea, and the cellular characteristics of the Asgard archaeal ancestor of eukaryotes.

## Acknowledgments

We thank F. Homa and V. de Jager for technical support and SURF (www.surf.nl) for the support in using the National Supercomputer Snellius.

## Funding

European Research Council Consolidator 817834 (TJGE)

Dutch Research Council VI.C.192.016 (TJGE)

Volkswagen Foundation 96725 (TJGE)

Simons Foundation (as apart of Moore-Simons Project on the Origin of the Eukaryotic Cell) 73592LPI; https://doi.org/10.46714/735925LPI) (TJGE, BJB)

SURF Cooperative grant no. EINF-2953. (TJGE)

Dutch Research Council VI.Veni.212.099 (JJEVH)

## Author contributions

Conceptualization: SK, TJGE

Data curation: SK, JJEVH, KP, KEA, VDA, DT

Orthology assignment: SK

Protein modeling: SK

Sequence homology searches: SK, JJEVH

Structural genomics analyses: SK, JJEVH

Genome data generation and curation: KEA, BJB, VDA

Phylogenetic analyses: SK, JJEVH, KP

Data interpretation: SK, JJEVH, KP, KEA, DT, TJGE

Funding acquisition: TJGE, JJEVH, BJB Supervision: TJGE

Writing – original draft: SK, JJEVH, TJGE

Writing – review & editing: SK, JJEVH, KP, DT, KEA, VDA, BJB, TJGE

## Data and materials availability

Custom code can be made available upon publication. All predicted structures, original multiple sequence alignments and IQ-TREE outputs will be made available upon publication. The uncollapsed phylogenies can be found on the iTOL (*61*) website: https://itol.embl.de/tree/62145192210399341699888333 (CINP, Figure 4A), https://itol.embl.de/tree/13722425212199811699868285 (COMMD, Figure 4C), https://itol.embl.de/tree/62145192210319901699902102 (Ufm1, Figure 4D).

## Supplementary Materials

Materials and Methods

Supplementary Text

Figs. S1 to S4

References (*62*–*107*)

Data S1 to S3

