## Supplementary Material for "Structure-based inference of eukaryotic complexity in Asgard archaea"

**Other Supplementary Materials for this manuscript include the following:**

Data S1 to S3

### Materials and Methods

Genome dataset selection

*Dataset assembly*

To construct a representative initial dataset, we retrieved all publicly available Asgard genomes from NCBI (*62*) up to October 6, 2022. This collection also included the recently published Asgard metagenome assembled genomes (MAGs) from the studies (*29*) and (*31*). To ensure data quality, MAGs were evaluated using CheckM v1.2.1 (*63*). Those MAGs with estimated completeness below 50% and estimated contamination exceeding 10% were identified as low quality and consequently excluded from the initial dataset. Taxonomic classification of the initial dataset was conducted using GTDB-Tk v2.3.2 (*64*) with default parameters. The final dataset comprised 936 genomes (Data SX) covering all known Asgard lineages. Gene prediction was performed using Prokka v1.14.6 (*65*) (options "--metagenome --kingdom Archaea").

*Phylogenomic inference*

To obtain an adequate outgroup dataset for inferring the phylogenetic relationships among the different Asgard lineages, we downloaded genus-level representatives of other archaeal lineages from the Genome Taxonomy Database (GTDB), release 214 (*66*). We based our selection on genome quality score (GQS), defined as GQS = completeness (%) - 5 x contamination (%), as described in (*67*). In cases where two genomes had equal GQS, a random selection was made between the two. The final outgroup dataset included 311 genus-level representatives classified as members of the Thermoproteota (excluding Korarchaeia, to avoid artefacts derived from their uncertain affiliation (*68*) and their strong thermophilic compositions (*13*)), Methanobacteria B, and Hadarchaeota lineages.

To infer the species tree, we performed phylogenomic analysis based on 47 non-ribosomal proteins, that were selected from a set of 200 markers previously identified as core archaeal proteins (Data S1) (*69*). Homologous sequences within the final genome dataset were recruited using PSI-BLAST (*70*) v2.10.0+ (-evalue 1e-10). All recruited sequences per taxon per protein marker were selected, aligned using MAFFT L-INS-i (*71*) v7.453, followed by trimming with trimAl (*72*) v1.4.rev22 (-gt 0.5) and removal of sequences with more than 60% gaps. We constructed the individual protein phylogenies using IQ-TREE (*73*) v2.1.3, incorporating model selection from ModelFinder (*74*). The best fitting model was selected among the combination of the LG, Q.pfam, and WAG models by adding the mixture model C20 with rate heterogeneity (+R4 or +G4) ("-mset LG+C20,Q.pfam+C20,WAG+C20 -mrate G4,R4 -mfreq ' ' "). We assessed branch robustness for each marker with 1000 ultrafast bootstraps (*75*) and SH-like approximate likelihood ratio tests (SH-aLRT) (*76*). From the resulting phylogenies, we removed sequences indicative of contamination, paralogy or horizontal gene transfer events and realigned and trimmed the remaining sequences as described above. The curated alignments were then concatenated into a supermatrix containing 1244 sequences. To mitigate effects related to compositional bias, we performed heterogeneous site removal using $\chi^2$-trimming (*77*) where the 50% most heterogeneous sites were removed resulting in an alignment of 8068 amino acid positions. We inferred a species phylogeny for the $\chi^2$-trimmed alignment using ModelFinder within IQ-TREE v2.1.3 to select among the LG+C10, Q.pfam+C10 and WAG+C10 models and rate heterogeneity components (+R4 or +G4). A PMSF approximation (*78*) of the best fitting model (WAG+C10+R4) using the resulting tree was then employed to reconstruct a final tree with 100 nonparametric bootstrap pseudo-replicates.

Clustering and selection of representative protein sequences

The dataset of 936 Asgard genomes comprised 2.68 million proteins. We assigned Asgard clusters of orthologous groups (AsCOGs) domains (*21*) to 2.1 million Asgard proteins according to the best hit to an AsCOG member using MMseqs2 (*79*) with '-e 0.001' and '-s 9' and at least 80% of the best hit had to be covered. Unassigned proteins (0.6 million) or protein fragments (0.2 million) of at least 60 amino acids were clustered *de novo* using MMseqs2 (*80*) v14.7e284 at 20% sequence identity and a coverage of 50%. We built sequence profiles for 14,467 (2,084,964 represented proteins) As-COGs and 22,846 (448,812 represented proteins) *de novo* clusters with at least 5 members. To select an evolutionary representative sequence per cluster, we searched members of the 37,313 clusters with at least five members against their respective cluster profile using MMseqs2 'mmseqs search', we ranked them based on their bit-score, and selected the highest-ranked sequence per cluster as the representative sequence (*79*).

Protein structure prediction

*Supplementing the ColabFold database with Asgard archaea proteins*

Protein structure prediction using AlphaFold2 (*25*) has been shown to generally perform poorer if few sequences can be aligned to the target sequence (*25*). We therefore wanted to evaluate whether adding our Asgard archaea protein dataset to the "genetic search" workflow of ColabFold (*32*), an accelerated adaptation of AlphaFold2, would increase overall prediction quality. To this end, implemented a version of the "genetic search" workflow of ColabFold that queries the Asgard protein dataset ("enriched") in addition to the default databases ("default"). For the enriched workflow, we added a third MMseqs2 sequence search step against the Asgard archaea protein database as after the searches against the two default ColabFold databases with the same parameters.

*Comparing performance of structure prediction algorithms for an Asgard archaeon*

To evaluate performance of different structure prediction algorithms as well as the ColabFold "default" versus the "enriched" database, we created a test set of 100 Asgard proteins (Data S1). We downloaded 100 randomly selected proteins of a reference Asgard archaeal proteome of *Candidatus* Prometheoarchaeum syntrophicum from UniProt (Data S1; Proteome ID: UP000321408; accessed on Jan 17, 2023). We first predicted structural models from the primary sequences using the protein language model based ESMfold v2.0.0 (*33*) with '-r 12'. To measure the quality of predictions we used the average predicted local difference test (pLDDT) score, ranging from low to high confidence (0-100). We considered predictions with an average pLDDT ≥ 80 as high-quality, as a compromise between the suggested pLDDT ≥ 90 for "high accuracy" and pLDDT ≥ 70 for "general correct backbone" according to (*25*). Secondly, we generated multiple sequence alignments with the "genetic search" module of ColabFold v1.3.0 (*32*) with default and enriched database, respectively. We then ran the prediction workflow of ColabFold on each alignment with the default "exhaustive" setting and with a premature stopping rule "early-stop" which is designed to save prediction time, i.e., the algorithm stops if either a pLDDT of at least 85 reached, or if the first prediction is below a pLDDT of 50 '--stop-at-score 85 --stop-at-score-below 50'. The "genetic search" module was ran on a computer with two AMD 7H12 64 cores 2.6Ghz 280W and 1 TiB of memory and the "prediction" module was run on a computer with four NVIDIA A100 40 GiB HBM2 memory.

*Protein structure prediction workflow*

Based on the highest ratio of high-quality proteins and lowest computational resource demands for our 100 test proteins, we opted for a hybrid approach of using protein language model and multiple sequence alignment-based prediction algorithms. We first used ESMfold v2.0.0 (*33*) with '-r 12' to calculate structural models for each representative Asgard archaeal protein. Secondly, structures with an average pLDDT < 80 in ESMfold were predicted again using ColabFold v1.3.0 (*32*) with the enriched database and the "early-stop" settings.

*Prediction of large proteins*

Large proteins that could not be folded with ESMfold v2.0.0 and ColabFold v1.3.0 because of exceeding memory demands, were attempted to be folded with ColabFold v1.5.2.

<u>Structure similarity searches</u>

*Best structural hit annotation*

We searched Asgard structures reciprocally against SwissProt predicted structures (downloaded July 8, 2022) using FoldSeek v6.29e2557 (*26*) 'foldseek search' with '--max-seqs 10000'. We retained the highest bit-score non-overlapping hits along the query sequence to accommodate fusion proteins and checked for reciprocal best hits. We mapped the annotation of the SwissProt best hits to each query protein. As described above, but unidirectionally, we searched Asgard structures against the PDB and UniProt50 databases (downloaded Feb 9, 2023).

*EggNOG annotation of SwissProt best hits*

Proteins representing the best SwissProt hits were mapped against EggNOG v5 (*81*) with the emapper user interface (http://eggnog-mapper.embl.de/) with default parameters and we extracted root NOG and eukaryotic NOG identifiers and functional categories.

*Identification eukaryotic hit enriched structures*

For each Asgard predicted structure we collected the best 10,000 hits of predicted UniProt50 structures (downloaded Feb 9, 2023). Per Asgard archaeal protein representative, we performed a one-tailed Fisher's exact test with the function 'fisher.test' and the 'alternative=less' parameter with Bonferroni correction with the function 'p.adjust' in R v4.2.1 (*82*) on the domain level taxonomy of hit UniProt proteins to test for a statistical enrichment in eukaryotic sequences. To test for eukaryotic enrichment in only the most similar proteins, we also performed the same statistical test using only the top 5% bit-score percentile of the hits. Structures with an enrichment in hits to eukaryotic proteins were classified as candidate isomorphic (i)ESPs, i.e., proteins that look structurally similar to proteins that are overrepresented in eukaryotes. We clustered all Asgard structures with Foldseek 'foldseek cluster' into clusters of isomorphic protein structures and identified structural clusters uniquely added with iESPs.

NCBI COG and KOG annotation of gene families

We created multiple sequence alignments for each Asgard archaeal protein cluster using FAMSA v2.2.2 (*83*) with '-refine_mode on'. We performed profile-profile searches with the HHsuite3 (*84*) program HHsearch v3.3.0 with parameters '-glob -M 50' against the profile COG-KOG database (ftp://ftp.tuebingen.mpg.de/pub/protevo/toolkit/databases/hhsuite_dbs/COG_KOG.tar.gz) (*85*).

*Mapping of ESPs described by Eme et al., 2023*

To identify conserved protein domains in the proteomes of the Asgard dataset, we used InterProScan v5.57-90.0 (*86*) with default parameters and using hidden markov models (HMM) from the databases AntiFam v7.0 (*87*), CDD v3.18 (*88*), Coils v2.2.1 (*89*), Gene3D v4.3.0 (*90*), MMobiDBLite v2.0 (*91*), PANTHER v15.0 (*92*), Pfam v35.0 (*93*), PIRSF v3.10 (*94*), PRINTS v42.0 (*95*), SFLD v4 (*96*), SMART v7.1 (*97*), SUPERFAMILY v1.75 (*98*), TIGRFAM v15.0 (*99*).

We then identified the AsCOG and de novo cluster protein domains containing at least 80% of the length of a Pfam or Interpro domains reported as ESPs (*13*).

Phylogenetic inferences

*iESP selection*

To illustrate how iESP confer information about the origins of eukaryotic functions and their proteins, we selected several iESPs for phylogenetic analysis, based on the following criteria: The Asgard archaeal query structure is well-covered (>80 % of protein length) by its alignment to its best structure hit; the best (eukaryotic) structure hit reciprocally has the Asgard archaeal query structure as its best hit; eukaryotic structures are overrepresented among the hits (Fig. 3B); the eukaryotic hit structures are consistent (are evidently homologous to one another); they comprise eukaryote-relevant functions; the query nor hit do not ostensibly embody very convoluted evolutionary histories, e.g., they, for example, do not consist of repeat domains or highly composite multidomain proteins; the Asgard archaeal query is unlikely to represent contamination, as it is found in more than one Asgard archaeal taxon. Finally, we demanded that the candidates did not have a well-scoring sequence-based hit (as derived by HHsearch) to eukaryotic sequences, hence they fall into the twilight zone of sequence homology (Figure 3C).

*Establishing remote sequence similarity between iESP and eukaryotic structure hits*

Subsequently, we established the iESPs, although divergent, do carry some signal in their sequences to link them to the eukaryotic proteins that they hit via structures. For this, we sought to gradually expand the homolog set of the iESP via manually supervised, iterative HMM searches. In each round, we checked the newly hit proteins before adding them to the multiple sequence alignment, as we ensured these are genuine homologs by inspecting both their sequences and (predicted) protein structures. We executed these profile HMM-based searches using online tools (HHpred, HMMer web server) as well as local hmmsearches onto our local databases (see description below). Note that, in addition to eukaryotic and Asgard archaeal sequences, we included bacterial and other archaeal sequences in the search database, since they might have homologs as well, and those homologs might in fact assist in connecting the iESP and related Asgard archaeal sequences to their eukaryotic structure hit.

*Selecting homologs for phylogenetic inference*

We made use of three sequence datasets for retrieving sequences for the phylogenetic analysis. First, we subsampled our in-house Asgard archaea set, including only a single representative protein set per species. This representative for a given species was selected based on the quality of the predicted proteomes, as reflected by their predicted completeness and contamination, measured by CheckM (*63*). Note that 'species' here signifies groups of genomes that can be clustered at the 95% average nucleotide identity (ANI) level. Second, we used a subsampled version of an in-house eukaryotic dataset (*100*), including 25 eukaryotic taxa of all of the major eukaryotic groups, taking the taxon with the best, most complete, predicted proteome quality, as measured by BUSCO (*101*). Third, we used a subsampled version of GTDB (r207) (*66*), of which first the Asgard archaea were removed, and then we selected the best assembly for each family, which was also based on the CheckM quality parameters. Using the final, most inclusive yet accurate profile HMM obtained, and our manually determined bitscore cutoffs (described above), we employed hmmsearch onto these three datasets and retrieved all sequences meeting the cutoff. Since COMMD and CINPL comprised virtually full-length hits, both at the structural comparisons as well as in our sequence similarity searches, we extracted the entire protein sequence of each hit protein. For Ufm1, we noticed that some hits in our sequence searches did not have a (near) full-length hit, and some had multiple hit regions, we only extracted the protein segment carrying the best hit region. For the major vault protein, in addition to the smaller full-length phylogeny (Figure 4A), we performed a broader phylogenetic analysis of the shoulder domain only, which is a type of Band 7 domain found in many prokaryotic and eukaryotic proteins (*44*, *102*), and which are united in the SPFH (for stomatins, prohibitins, flotillins and HflK/C) family 'clan' (https://www.ebi.ac.uk/interpro/set/pfam/CL0433/entry/pfam/).

*Phylogenetic analysis and annotation of the phylogeny*

For each family, we choose inferred gene trees using multiple sequences alignments generated by MAFFT (v7.505, mode L-INS-i) (*71*) and the web server of PROMALS3D (*103*). For the latter, we used the default options, except for detecting and using homologs with 3D structures (included DaliLite v5 (*104*)), pairwise alignments between input 3D structures (included DaliLite) and aligning sequences within groups in the first alignment stage (PROMALS instead of MAFFT). We supplemented PROMALS3D with predicted protein structures from diverse sequences from the AlphaFold Protein Structure Database, as well as available structures from our own structure prediction (described above), i.e. of the iESP and, if available, other Asgard archaeal homologs. Before inferring the gene trees, we trimmed the multiple sequence alignment using BMGE (v1.12, settings: -m BLOSUM30 –h 0.6 -g 0.7 -b 3) (*105*), which selects good-quality aligned positions. However, in some cases (e.g., COMMD MAFFT alignment), this would result in a very short alignment, due to which we switched to trimAl (v1.4.1, mode 'gappyout') (*72*). For phylogenetic inference in a maximum-likelihood framework, we used IQ-TREE (v.2.0.3, settings -B 1000 -m MFP -mset LG,JTT,Q.pfam,WAG,LG+C20,LG+C40,LG+C60,LG+C20+R+F,LG+C40+R+F,LG+C60+R+F,WAG+C20,WAG+C40,WAG+C60,WAG+C20+R+F,WAG+C40+R+F,WAG+C60+R+F,JTT+C20,JTT+C40,JTT+C60,JTT+C20+R+F,JTT+C40+R+F,JTT+C60+R+F,Q.pfam+C20,Q.pfam+C40,Q.pfam+C60,Q.pfam+C20+R+F,Q.pfam+C40+R+F,Q.pfam+C60+R+F) (*73*) to first select the best evolutionary model using ModelFinder (*74*) and then infer a phylogeny with 1000 ultrafast bootstraps (*75*). For each iESP/family, we subsequently selected the phylogeny displaying the most informative and probably accurate tree, which entailed post-hoc selecting the alignment algorithm (MAFFT-L-INS-i versus PROMALS3D) (based on ultrafast bootstrap support values at key branches, and monophyly of expected monophyletic sequence groups). We coloured the branches in the tree according to the species group the sequences belong to: Eukaryota, Asgard archaea, Archaea (other) and Bacteria. We also annotated the eukaryotic clades with the names of their proteins, specifically labelling each clade reflecting a single gene in the last eukaryotic common ancestor (LECA). Trees were visualized using iTOL (*61*).

*Visual representation of protein structures*

Structural models were either visualized in ChimeraX v1.6.1 (Fig. 4B-F) (*106*) or in R with the 'r3dmol' package v0.1.2 (Fig 4G-H) (https://github.com/swsoyee/r3dmol) (*107*).

Data availability

The uncollapsed phylogenies of Fig. 4E, 4G, 4H, S4A can be found on the iTOL website: https://itol.embl.de/tree/13722425212199811699868285 (COMMD, Fig. 4E), https://itol.embl.de/tree/62145192210319901699902102 (Ubiquitin fold modifier 1, Fig. 4G), https://itol.embl.de/tree/62145192210399341699888333 (CINP, Fig. 4H), https://itol.embl.de/tree/1372242521225521718619040 (MVP shoulder, Fig. S4A).

### Supplementary Figures

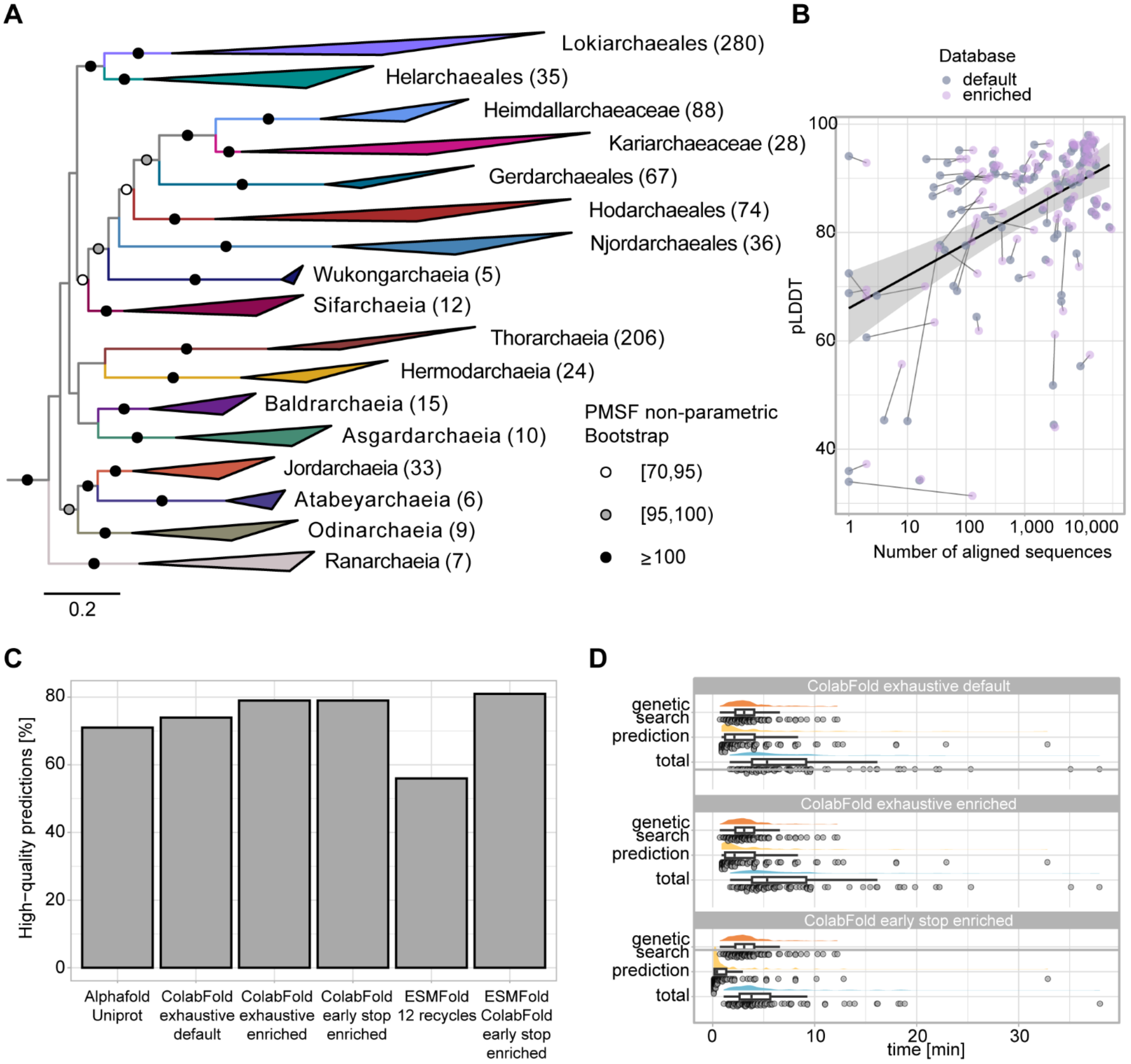


**Fig. S1. Asgard phylogenomic tree and comparison of structural prediction algorithms** (**A**) Maximum-likelihood phylogenetic tree of 935 Asgard archaeal genomes, using Euryarchaeota and Thermoproteota archaeal representatives as outgroup (not shown). The tree is based on 47 concatenated non-ribosomal proteins (8068 sites and 1244 taxa), using IQ-TREE under the WAG+C10+R4 model. PMSF approximated non-parametric bootstrap support ≥70 is indicated on branches. Scalebar represents the average expected substitutions per site. (**B-D**) Evaluation of different aspects of the structure prediction workflow in the following panels were performed on a set of 100 proteins of '*Candidatus* Prometheoarchaeum syntrophicum' (see Methods). (**B**) Number of aligned reference sequences (x-axis) and average structure model pLDDT (y-axis) with default (blue) and enriched (purple) database. (**C**) Number of high-quality structure predictions (pLDDT ≥80) based on different predictions strategies. (**D**) Inference times of ColabFold prediction modules with different inference strategies, including the default setting and database, or the enriched database with either default settings or an early stop criterion (see Methods).

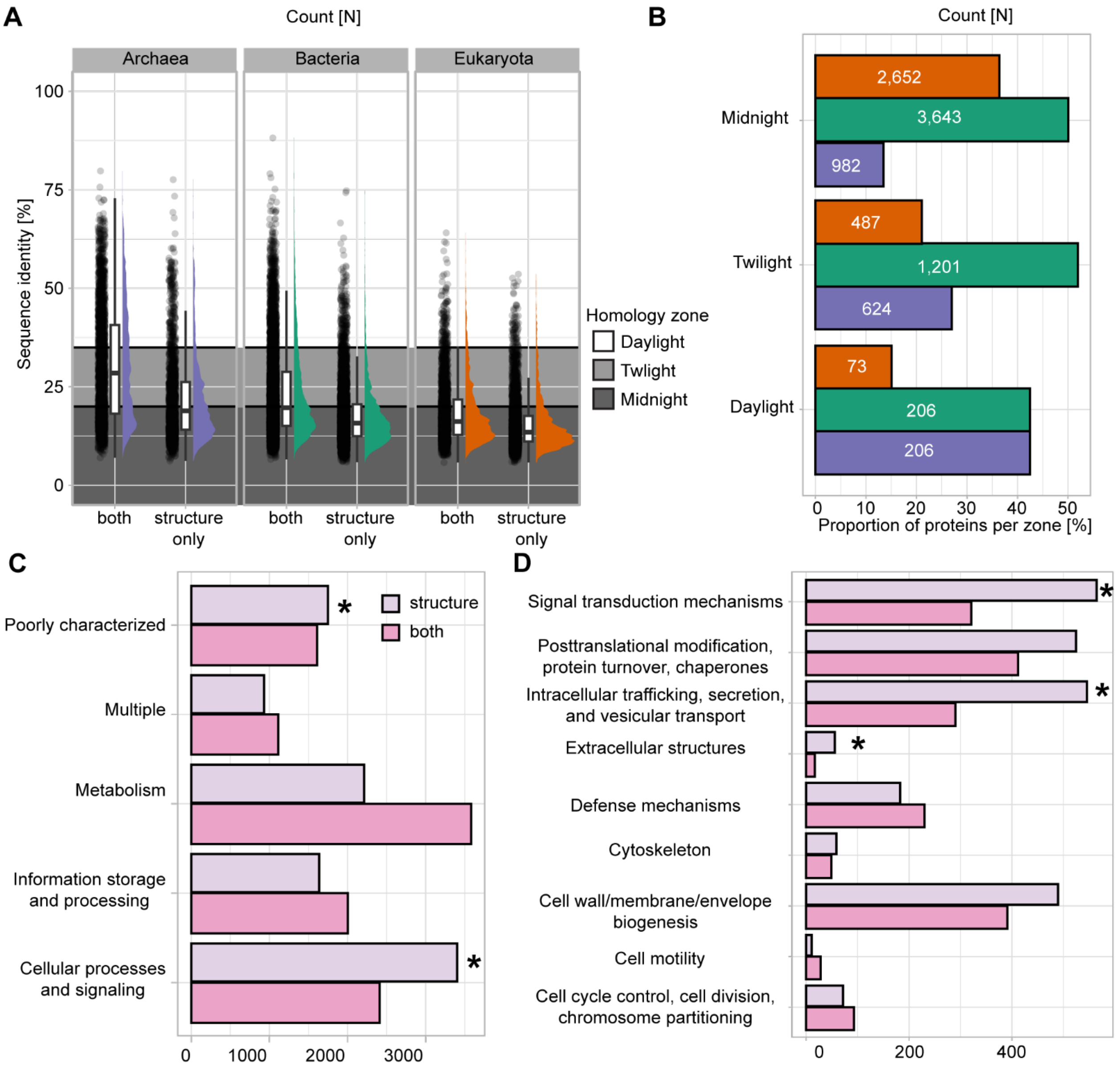


**Fig. S2. Comparative assessment of Asgard archaeal structural clusters. (A)** Sequence identity of representative Asgard archaeal proteins to SwissProt best-hit protein, segregated by the domain of life. **(B)** Proportion of proteins per domain (colors follow panel A) per sequence homology zone. **(C)** Bar plot indicating the number of hits per functional category of the best Swiss-Prot hit per Asgard archaeal representative protein. Asterisks indicate a functional enrichment based on a one-tailed Fisher's exact test with Bonferroni correction of "Cellular processes and signaling" and "Poorly characterized" with $9.3 \cdot 10^{-77}$ and $1.4 \cdot 10^{-7}$, respectively. **(D)** Bar plot indicating the number of hits per functional subcategory within "Cellular processes and signaling". Asterisks indicate a functional enrichment based on a one-tailed Fisher's exact test with Bonferroni correction of "Intracellular trafficking, secretion, vesicular transport", "signal transduction", and "extracellular structures" with the respective adjusted p-values: $5 \cdot 10^{-6}$, $3 \cdot 10^{-4}$, $6.3 \cdot 10^{-4}$.

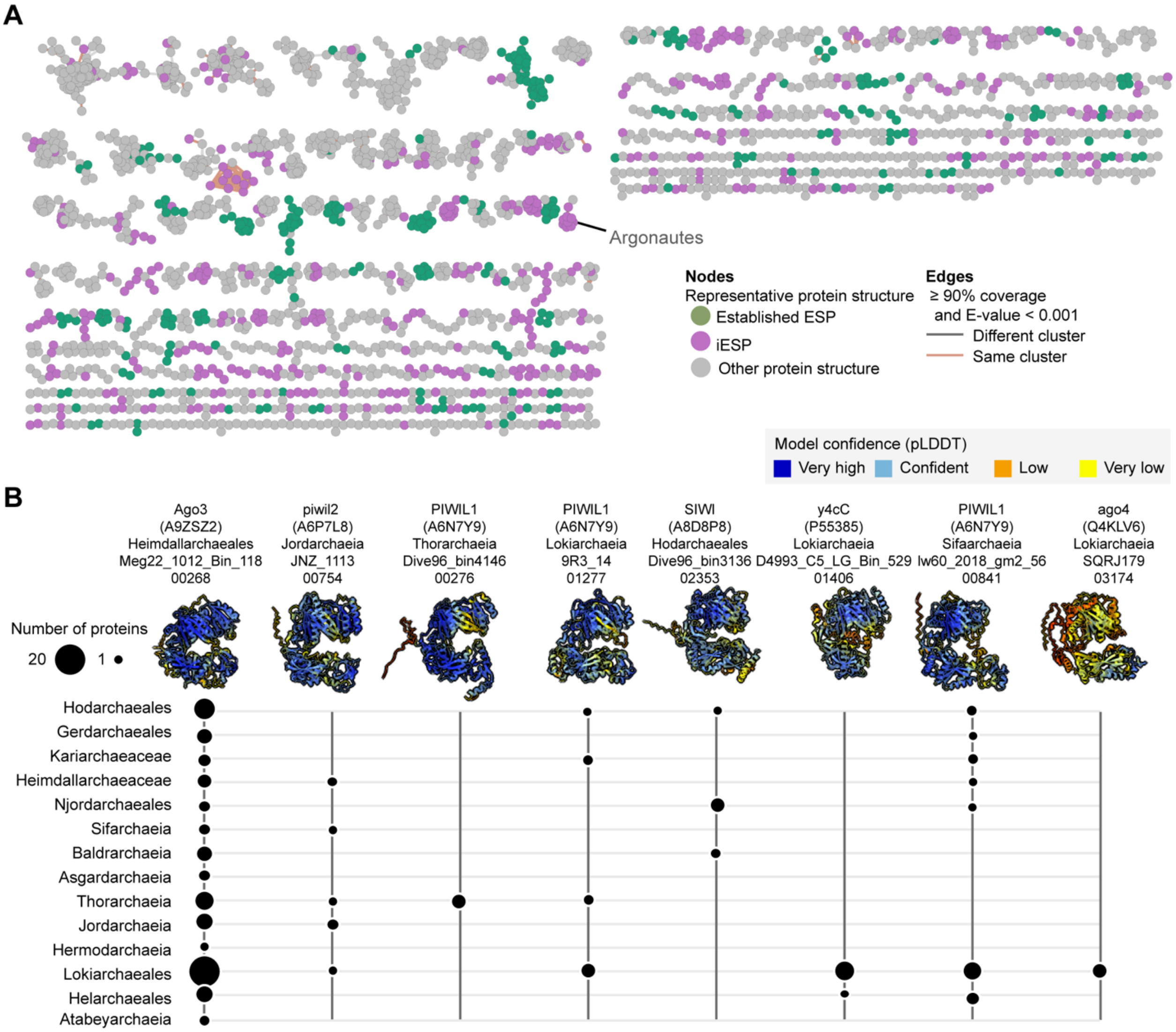


**Fig. S3. Analyses of the Asgard archaeal protein structure similarity network.** (**A**) Subgraph complementing the protein structure similarity network depicted in Figure 3, once again highlighting Argonaute proteins. (**B**) Distribution across Asgard archaeal groups of eight Asgard archaeal Argonaute-related iESPs contained in a single structural cluster.

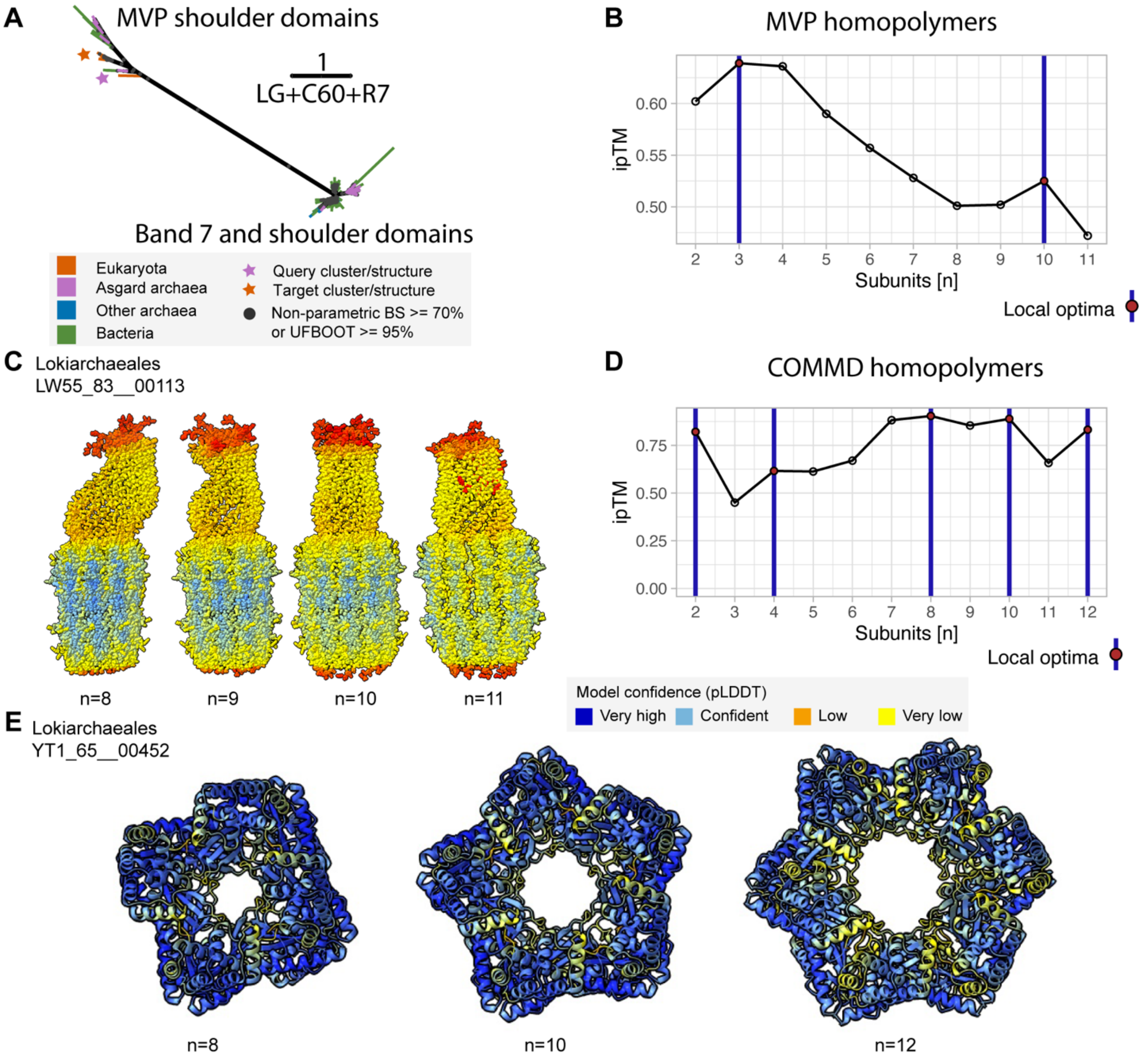


**Fig. S4. Phylogenetic and structural analyses of iESPs. (A)** Protein domain phylogeny based on Band 7, MVP and related shoulder domains. The depicted phylogenetic tree is based on 90 aligned positions and was generated under the LG+C60+R7 model (see Methods). **(B)** ipTM score of Asgard archaeal MVP homopolymers modeled with different numbers of subunits with local optima highlighted. **(C)** Multimer model of Lokiarchaeial MVP with different number of subunits. **(D)** ipTM score of Asgard archaeal COMMD homopolymers modeled with different numbers of subunits with local optima highlighted. **(E)** Homo-multimer model of Lokiarchaeial COMMD-containing protein with different number of subunits.

### Supplementary Data

#### Data S1. (separate file)

Spread sheet with genome information of the outgroup genomes used for Fig. S1A and the dataset of 936 Asgard archaeal draft genomes.

#### Data S2. (separate file)

Spread sheet of UniProt protein IDs and annotations of sampled *Candidatus* Prometheoarchaeum syntrophicum.

#### Data S3. (separate file)

Spread sheet including the annotation of structures in the of ESPs and iESP structural clusters.
